# Risk Factors of Mortality of Hospitalised Adult Burn Patients a Malawian Tertiary Hospital Burns Unit

**DOI:** 10.1101/421982

**Authors:** Stephen Kasenda, Donnie Mategula, Geoffrey Elihu Manda, Tilinde Keith Chokotho

## Abstract

**Introduction:** Malawi has the highest rates of mortality directly or indirectly associated with burn injuries in Southern Africa. There is however no published literature on risk factors of mortality among adult patients.

**Methods:** We conducted a retrospective cross sectional audit records of patients admitted at the burns unit of Queen Elizabeth Central Hospital (QECH) between the years 2007 and 2017. Death due to burns was our outcome of interest. We collected patient data including demographic information, details of the burn injury and its management and determined how these factors were associated with the risk of death using Person Chi square tests in a univariate analysis and likelihood ratio tests in a multivariate logistic regression model. We also determined the odds ratios of death within the categories of the risk factors after adjusting for important variables using a logistic regression model.

**Results:** An analysis of 500 burns patient records showed that 132(26.4%) died during the 10-year period. The lethal area for 50% of burns (LA50) was 28.75% and mortality reached 100% at 40% total burn surface area. The following variables were found to be significantly associated with mortality after controlling for confounders: scalds (OR 0.13; 95% CI 0.05-0.33; <0.0001), increasing total burn surface area (p<0.0001), time lapse to hospital presentation between 48 hours and one week(OR 0.27; 95%CI 0.11-0.68; <0.0001), inhalation burns (OR 5.2; 95% CI 2.0-13.3 p 0.0004) and length of hospital stay greater than two months (OR 0.04 95, CI 0.01-0.15; P<0.0001).

**Conclusions:** Risk factors for mortality are connected by their association with post-burn hypermetabolism. Further studies to are needed to identify the best and cost-effective ways of preventing death in burn patients.

## INTRODUCTION

A burn is defined as an injury to the skin or other organic tissue primarily caused by heat or due to radiation, radioactivity, electricity, friction or contact with chemicals.(1)It is estimated that 90% of the worldwide burns associated mortality are in the low and middle income countries with Africa alone accounting for 15% of burns mortality(2,3) A systematic review of scientific papers from 14 African countries showed that Malawi has the highest burns related mortality in Southern Africa (22%) which is also higher than the average burn mortality of Africa (17%) for all age groups.(4) Despite the estimated significant burden of disease, there is paucity of data on the quality of burn care and outcomes in Malawi and Sub-Saharan Africa in general. There is also a lack of established mechanisms to reduce burn related mortality and morbidity(2) We undertook, to the best of our knowledge, the first ever retrospective study of burns among adult patients from 17 years old and above admitted in the Queen Elizabeth Central Hospital (QECH) burns unit from 1^st^ June 2007 to 31^st^ May 2017 with the aim of determining the prevalence and risk factors of mortality among them.

## METHODS

### Study design and setting

The study was a retrospective cross sectional audit of patient records at Queen Elizabeth Central Hospital (QECH) which is the largest referral (tertiary) hospital in Malawi with a bed capacity of over 1000. The facility also treats patients referred directly from primary level health facilities due to absence of a secondary level hospital in Blantyre district where it is located. QECH is also the location one of the two burns units in the country with a bed capacity of 32.

### Study population

All patients aged 17 years and above with any type of burn injury admitted in the QECH Burns Unit between 1^st^ June 2007 and May 2017 met the inclusion criteria of this audit. Patients excluded from the study were those who; had no documentation beyond the admission process, were pronounced dead on arrival, presented for the first time with complications of burns and those with non-burn-related issues.

### Data collection

We searched the surgery department electronic records as well as the ward registers in order to identify burn patients of at least 17 years age upon admission who were admitted in the burns unit between 1^st^ June 2007 and May 2017. The following type of data was extracted from patient files; demographic data (admission date, discharge date, age, gender, referring facility/residence); injury-related data (place of injury, time to hospital presentation, first aid before presentation, aetiology of burn injuries, depth and Total burn surface area, circumstances of the burn, presence of inhalation burns, presence of comorbidities and quality of control of comorbidities); initial and subsequent management (fluid management, physiotherapy and nutritional support); presence of fever and patient outcomes.

### Data entry and analysis

Data entry and initial cleaning was done using Microsoft Excel 2007 and the final data cleaning and analysis was done using Stata Cooperation version 15. The outcome variable of death was categorised to a binary set of either death or alive. Patients that absconded or were transferred to other facilities or burns clinic were assumed to be alive. The exploratory variables to be explored as risk factors of death were categorised into meaningful categories that have a scientific backing where possible and in such a way that data sparsity in terms of the outcome of death was avoided. The initial univariate analysis was done by cross tabulation of the potential risk factors with the outcome of death, taking note of the Pearson chi-square p values of the association with the outcome of death. Odds ratios were calculated, comparing the odds of death in a category of the variable to a selected baseline category of the same risk factor while taking note of the 95% confidence interval. With reference from previously published studies(5) and expert opinion, we developed causal framework of causes death in light of the variables we collected as a Directed Acyclic Graph (Figure 1) using the software DAGitty version 2.3. We independently assigned variables of interest as exposure variables with respect to the outcome of death and then identified variables that need to be adjusted for. A logistic regression model was used to calculate the adjusted odds ratios of death within the categories of a potential risk factor taking note of 95% confidence intervals and the likelihood ratio p values of the association of the variable with the outcome of death after adjusting for variables identified in our causal framework.

**Figure 1:**
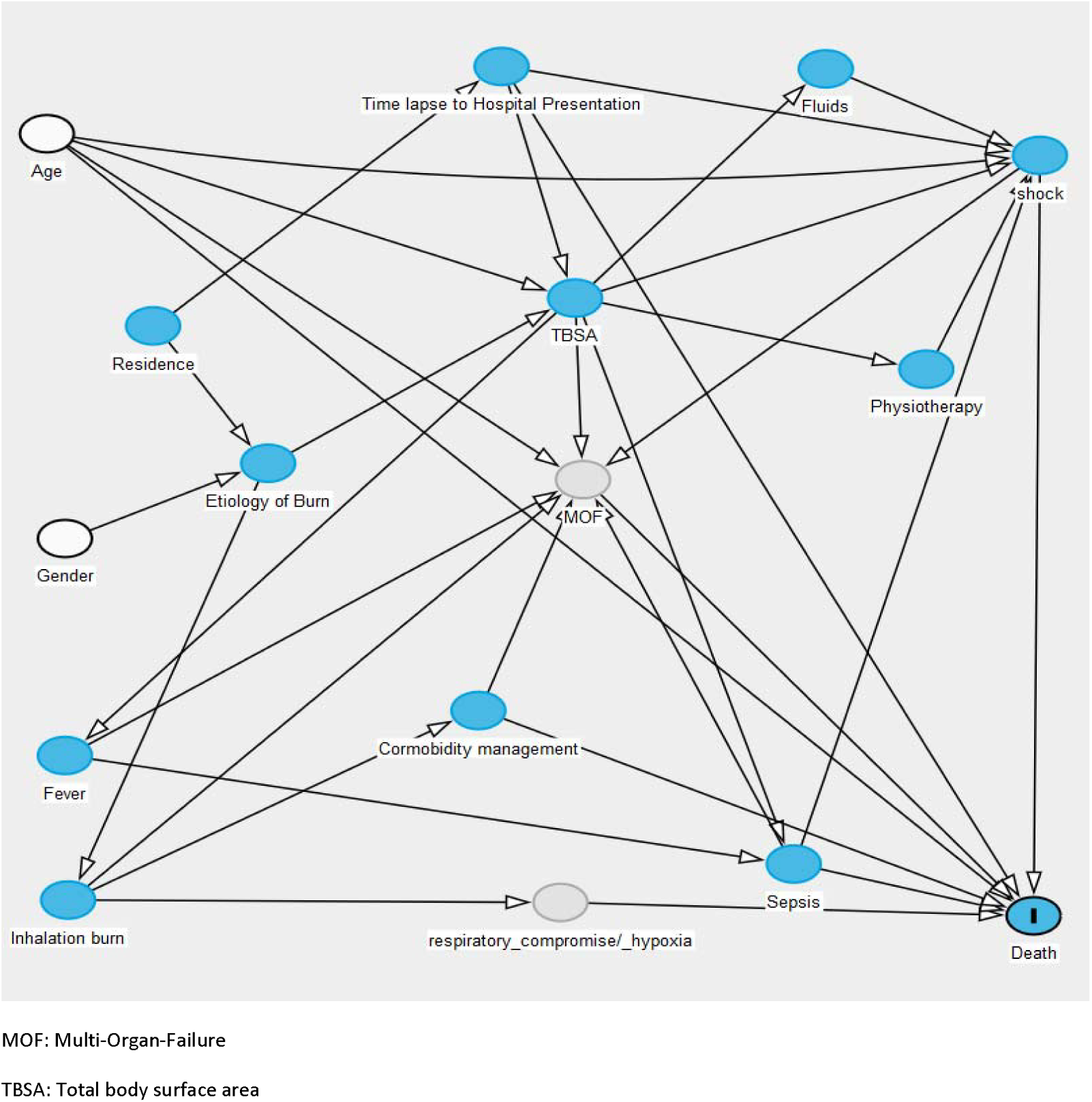
Directed Acyclic Graph for risk factors of death in burns patients MOF: Multi-Organ-Failure TBSA: Total body surface area

### Ethical approval

Ethical approval was obtained from the College of Medicine Research and Ethics Committee (COMREC); COMREC reference number P.09/17/2275.

## RESULTS

Out of the 779 names obtained from the patient registers in the unit, 500 patient files met the inclusion criteria, had their outcomes recorded and were available at the time of file retrieval. The median age at the time of presentation was 32 (IQR: 25-45; range 17-92 years) and 260 (52%) of the patients were male. A majority of the patients (57.6%) were 18-35 years old seconded by those who were 36-53 years old (23.4%). The rest of the patient age categories contributed 10% or less each. Three hundred and fifty six (71.2%) of the patients were from rural areas and the rest from urban areas. Most of the burns were: domestic (73.6%), accidental (52.8%) and of flame aetiology (78.4%).

An outcome of death was recorded for 132(26.4%) of the patients with 29(22%) of all deaths occurring within days of admission and 68(50.7%) deaths occurring within the first week of admission. Out of the 366 patients that were recorded to be alive upon discharge, 12 were transferred to a secondary level hospital, one transferred to a South African hospital and 10 were transferred to the outpatient burns clinic. The specific causes of death were recorded in 30 patient files only based on the clinical circumstances surrounding the death and not autopsy findings. Causes of death as recorded in the patient files were: infections (36.7%), severe burns (26.7%), shock (20%), respiratory failure (6.7%), inhalation injury (6.7%) and aspiration (3.3%).

According to the univariate analysis done (table 1) on all the categorical variables tested, the following factors were strongly associated with mortality(P values in the table 1): age, gender, residence, burn aetiology, time to hospital (QECH) presentation, Alcohol intoxication at the time of burn, circumstances of the burn, inhalation burns, poorly controlled co-morbidities, total burn surface area, burn depth, *fluid resuscitation*, type of wound management, length of hospital stay, and presence of fever. The place of burn however had no statistically significant effect on mortality.

**Table 1:** Various factors’ crude odds ratios (OR) of death and the chi square p value of association with death

| VARIABLE | CATEGORY | SIZE (N) | DEATHS (%) | OR (95% CI) | PEARSON $\chi^2$ P VALUE |
| --- | --- | --- | --- | --- | --- |
| **Gender** | Female | 240 | 57(21.9) | 1 | 0.018 |
|  | Male | 260 | 75(31.3) | 0.62 (0.41-0.92) |  |
| **Age** | <18 years | 20 | 2(10) | 1 | <0.001 |
|  | 18-36 years | 288 | 62(21.5) | 2.5 (0.6-11.0) |  |
|  | 37-52 years | 117 | 30(25.6) | 3.10 ( 0.67-14.44) |  |
|  | 53-72 years | 50 | 23(46.0) | 7.67 ( 1.44- 40.88) |  |
|  | >72 years | 23 | 15(65.2) | 16.88 ( 2.12-134.37) |  |
| **Aetiology[1]\*** | Flame | 392 | 127(32.4) | 1 | <0.001 |
|  | Scald | 87 | 5(5.8) | 0.13 (0.05-0.33) |  |
| **Place of Burn** | Home | 368 | 103(28.0) | 1 | 0.251 |
|  | Work | 56 | 11(19.6) | 0.63 (0.31- 1.27) |  |
|  | Other | 32 | 6(18.80) | 0.59 ( 0.24-1.49) |  |
| **Circumstance of Burn** | Accidental | 264 | 76(28.8) | 1 | 0.002 |
|  | Alcohol intoxication | 17 | 10(58.8) | 3.53 (1.28-9.76) |  |
|  | Seizure | 153 | 29(18.95) | 0.58 ( 0.36-0.94) |  |
|  | Intentional | 24 | 5(20.8) | 0.65 (0.23-1.81) |  |
| **Inhalational burns** | None | 475 | 116(24.4) | 1 | <0.001 |
|  | Present | 25 | 16(64.0) | 5.50 ( 2.33- 13.01) |  |
| **TBSA\*\*** | <10% | 198 | 10(5.1) | 1 | <0.001 |
|  | 10-20% | 124 | 29(23.4) | 5.74 (2.60-12.66) |  |
|  | 21-29% | 54 | 27(50.0) | 18.80 (7.09- 49.82) |  |
|  | 30-39% | 38 | 20(52.63) | 20.89 (7.24- 60.28) |  |
|  | ≥40% | 43 | 42(97.7) | 789.60 (12.85- 4.85e+04) |  |
| **Management type** | Conservative | 306 | 119(38.9) | 1 | <0.001 |
|  | Surgery | 194 | 13(6.7) | 0.11 (0.06-0.22) |  |
| **Comorbidities** | No comorbidities | 284 | 83(29.2) | 1 | 0.031 |
|  | Unknown | 160 | 30(18.8) | 1.79 ( 1.11- 2.88) |  |
|  | Well controlled | 2 | 0(0.0) | 0 |  |
|  | Poorly controlled | 54 | 19(35.2) | 2.35 ( 1.17- 4.72) |  |
| **Fever** | No | 187 | 35(18.7) | 1 | 0.003 |
|  | Yes | 249 | 78(31.3) | 1.98 (1.25- 3.14) |  |
| **Residence** | Rural | 364 | 118(32.4) | 1 | <0.001 |
|  | Urban | 136 | 14(10.30) | 0.24 (0.13- 0.44) |  |
| **Fluids indicated** | No | 249 | 26(10.4) | 1 | <0.001 |
|  | Yes | 209 | 95(45.5) | 7.15 (4.19-12.18) |  |
| **Fluids given** | No | 380 | 57(15.0) | 1 | <0.001 |
|  | Yes | 109 | 70.(64.2) | 10.17 ( 5.89-17.56) |  |
| **Physiotherapy needed** | No | 142 | 16(11.3) | 1 | <0.001 |
|  | Yes | 324 | 111(34.3) | 4.10 (2.28-7.37) |  |
| **Physiotherapy done** | No | 415 | 119(28.7) | 1 | 0.008 |
|  | Yes | 82 | 12(14.6) | 0.43 (0.22-0.82) |  |
| **Nutritional Support needed** | No | 268 | 20(7.5) | 1 | <0.001 |
|  | Yes | 198 | 108(54.6) | 14.88 (7.95-27.86) |  |
| **Nutritional Support done** | No | 489 | 128(26.2) | 1 | 0.779 |
|  | Yes | 3 | 1(33.3) | 1.41 (0.13-15.73) |  |
| **Time lapse to presentation** | <8 hours | 37 | 19(51.4) | 1 | <0.001 |
|  | 8- 24 hours | 110 | 37(33.6) | 0.48 (0.22- 1.036) |  |
|  | >24-48 hours | 112 | 33(29.46) | 0.40 (0.18- 0.86) |  |
|  | >48- 1 wk | 111 | 18(16.22) | 0.18 (0.08- 0.44) |  |
|  | >1 wk | 118 | 24(20.34) | 0.24 (0.11- 0.55) |  |
| **Length of hospital stay** | <1 wk | 113 | 68(60.2) | 1 | <0.0011 |
|  | 1wk -1 month | 192 | 35(18.23) | 0.15 ( 0.08- 0.26) |  |
|  | 1-2 months | 70 | 12(17.4) | 0.14 (0.06- 0.31) |  |
|  | >2 months | 125 | 17(13.6) | 0.10 (0.05- 0.22) |  |

[1] ETIOLOGY: other burn aetiologies not included in this table did not result in mortality

The median total burn surface area was 12% (IQR 5.5% - 21.5%). One hundred and ninety eight (39.6%) patients had TBSA <10% and mortality rate was least prevalent in among these. The odds of death increased with each subsequent category with an exponential increase occurring in the >40% category. Burn surface area that resulted in 50% mortality (LA50) was 28.75% and 100% mortality occurred at 40% TBSA. The odds of death also increased with each TBSA category with an exponential increment occurring in the >40% TBSA group. There was a mean difference of 7.9% between patients that presented within 48 hours post-burn (median 13.5%; mean 21.8±1.4%) and those that presented later (median 9.5%; mean 13.6±0.9%).

One hundred and ninety four (38.8%) patient had surgical intervention during their admission with median time to surgery was 24.5 days (IQR 11-44.8 days). According to the 188 recorded outcomes surgical intervention resulted in 90.9% less odd of death when compared to conservative management. Among those who had surgery and recorded outcomes, only 15 (8%) surgical intervention within 48 hours. The discrepancy in the sample sizes of the two surgical intervention categories made it difficult to accurately assess the impact of early surgery compared to delayed surgery. Excision of infected wounds was done only twice during the whole study period.

Further analysis was done on fluid management, physiotherapy and nutritional support and their effect on mortality comparing occasions where they were indicated or not to when they were actually done or not.(Table 2) unnecessary use and lack of fluid administration when indicated was recorded in 7(1.4%) and 104 (20.8%) of patients respectively. Administration of intravenous fluids in occasions where they were indicated was strongly associated with a 6.8 (CI 3.1-13.7, P<0.0001) greater odds of death when compared to those who had a similar need but did not get fluids. Institutional nutritional support was offered to three (0.6%) patients only in throughout the entire study period and there was no evidence of increased or reduced odds of death when indicated and given compared to when indicated and not given (OR 0.4; CI 0.03-4.5; P value 0.4387). Administration of physiotherapy was associated with 0.27 less odds of mortality among those for whom it was indicated (CI 0.135-0.532; P value= 0.0001). No other supportive management measures were availed to the patients apart from physiotherapy, nutritional support and fluid management.

**Table 2:** Indication vs. actual administration of IV fluids, nutritional support and physiotherapy.

|  |  | IV Fluids indicated |  |  |  |
| --- | --- | --- | --- | --- | --- |
|  |  | No |  | Yes* |  |
|  |  | Alive | Died | Alive | Died |
| IV Fluids given | No | 216 | 25 | 79 | 25 |
|  | Yes | 6 | 1 | 31 | 67 |

|  |  | Nutritional support needed |  |  |  |
| --- | --- | --- | --- | --- | --- |
|  |  | No |  | Yes** |  |
|  |  | Alive | Died | Alive | Died |
| Nutritional support done | No | 248 | 20 | 84 | 106 |
|  | Yes | 0 | 0 | 2 | 1 |

|  |  | Physiotherapy support needed |  |  |  |
| --- | --- | --- | --- | --- | --- |
|  |  | No |  | Yes*** |  |
|  |  | Alive | Died | Alive | Died |
| Physiotherapy support done | No | 124 | 16 | 146 | 99 |
|  | Yes | 1 | 0 | 66 | 12 |

\*OR 6.8(95% CI 3.4 - 13.7) p value <0.0001
\*\* OR 0.4(95% CI 0.03 -4.5) P value 0.4387
\*\*\* OR 0.27(95% CI 0.135222 -0.531696) P value =0.0001

Table 3 shows the logistic regression odds ratios after adjusting for variables determined through a causal framework developed by the authors. After adjusting for age, gender residence and time lapse to presentation, there was strong evidence that the aetiology of the burn was strongly associated with death (p<0.0001). Those who had a scald had a 0.13(95% CI 0.05 -0 .33) lower odds of death compared to those who had a flame caused burn. The total burn surface area (TBSA) was also strongly associated with the risk of death ((p<0.0001) with those who had a TBSA of greater than 40% have a 783 greater odds of death compared to those who had a TBSA of <10% (Adjusted OR 782.07 95% CI 89.9-6801.1). If a patient had an inhalation burn, he or she had a nearly 5 times increased risk of dying compared to if they had not.(adjusted OR 5.2 95%CI 2.0 - 13.3 P value 0.0004).

**Table 3:**
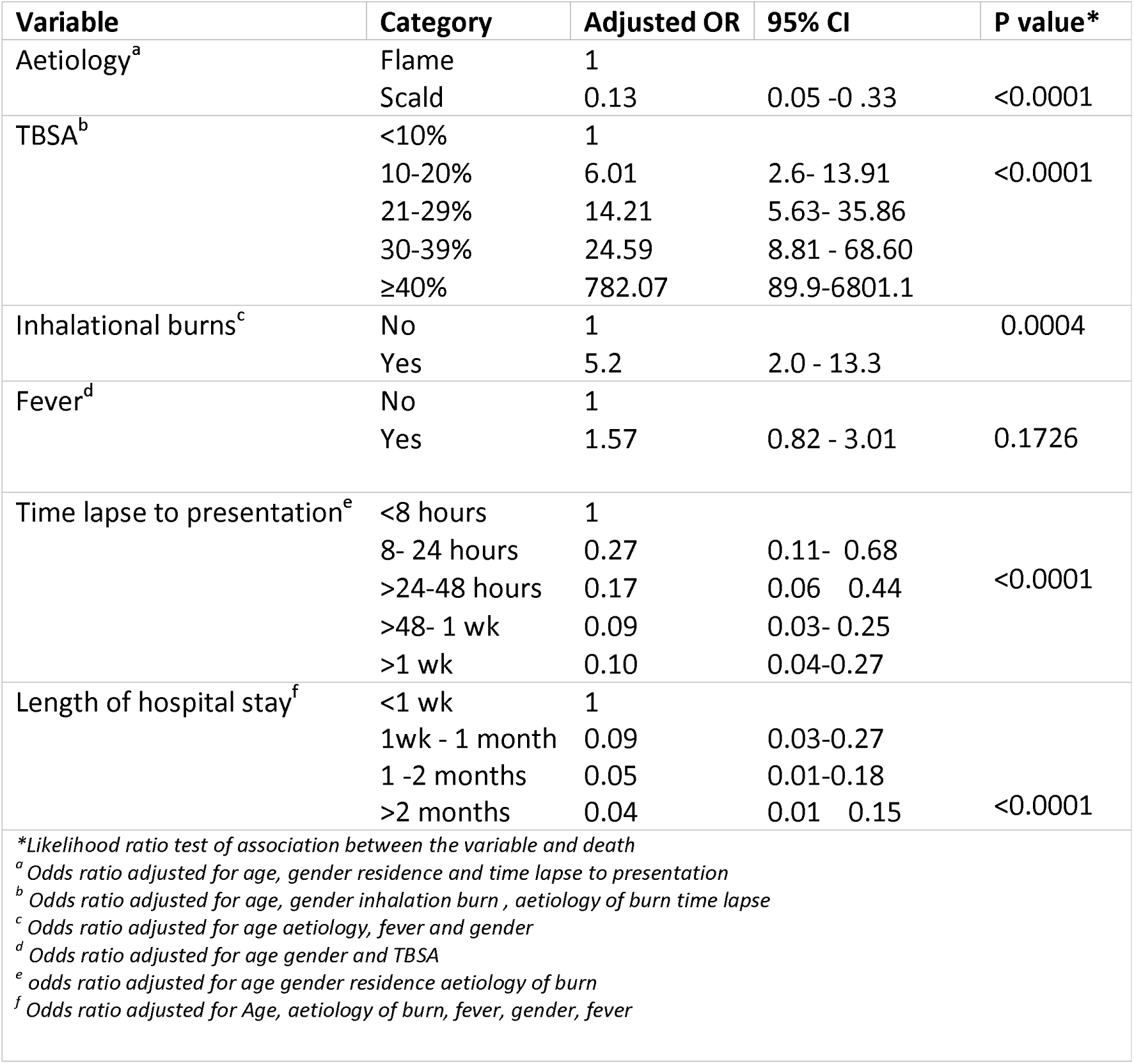
Logistic regression adjusted odds ratios for the association of selected variables with death.

The time lapse to presentation and the length of hospital stay were also strongly associated with the risk of mortality (p value for each <0.0001). Those that presented to the hospital within 8-24 hours had a 73% reduced risk of death compared to those that presented less than 8 hours of the burn.(adjusted OR 0.27 95% CI 0.11-0.68). Those that presented to the hospital more than 48 hours to 1 week had a 91% reduced risk of death compared the same group as above (adjusted OR 0.09 95% CI 0.03-0.25). Compared to those that stayed in the hospital for less than one week, those that stayed more than one week to one month had 91% reduced risk of death (adjusted OR 0.09 95% CI 0.03-0.27).Similarly, those that stayed for more than 2 months had a 96% reduced risk of death compared to those that that stayed less than a week (Adjusted OR 0.04 95% CI 0.01 -0.15).

## DISCUSSION

In this study, we reviewed records of 500 patients admitted at QECH between 2007 and 2017 of which 132(26.4%) resulted in death during the time of admission. We have determined that age, gender, burn aetiology, total burn surface area, burn depth, presence of inhalation burns, circumstances of the burn, quality of co-morbidity management, type of management for the burn wounds, patients residence, time to hospital presentation, length of hospital stay and supportive as well as resuscitative management administration are the risk factors that are strongly associated with mortality. Using a direct acyclic graph we identified burn aetiology, total burn surface area, inhalation burns, time to hospital presentation, length of hospital stay and presence of fever as the minimum set of exposure variables that (individually or combined in different ways) may lead to death. These results are consistent with results from international studies which found the same factors to be associated with mortality.(6–9) These results also confirm the existence of high burn related mortality prevalence Malawi.(4,6)

A comparison of our results with those of previous studies makes it evident that the factors associated with burns related mortality in our setting are predominantly linked by their association with post-burn hypermetabolism.(5,10–13) Hypermetabolism has been described as a major contributor (both direct and indirect) to the overall burns related mortality and morbidity especially in the Flow phase of a burn which starts after at least two (2-5) days following a burn and lasts up to 36 months.(12,14,15) The association between hypermetabolism and mortality is more evident with our findings of LA50 at 28.75% TBSA and 100% mortality at 40%TBSA since the post-burn metabolic rate increment is 50% and 100% at TBSA’s of 25% and 40% respectively.(16)

The magnitude and duration of this phase is determined by increased levels of catecholamines, glucagon, and cortisol which induce increased proteolysis, lipolysis, glycolysis.(5,11,12,14,15,17) The wound size and time to excision also contribute to hypermetabolic response by releasing pro-inflammatory mediators and cytokines and by attracting neutrophils(17) These changes have been associated with glucose intolerance, reduced oxygen consumption, raised body temperature (by an average of 2^11^C), immunosuppression, increased cardiac output and when in excess can lead to cardiomyopathy, focal necrosis, multi organ failure and eventually death.(14,17) Based on our understanding of the mechanisms that drive hypermetabolism, it is evident that hypermetabolism and its effects can be alleviated by implementation of interventions (with locally available resources) that limit or inhibit the pathways involved in the pathological cascade. These interventions include control of: pro-inflammatory cytokines^1^, catecholamine release, catabolic pathways and hyperglycemia

Catecholamines are the primary mediators of the hypermetabolic response and blockade of its effects has proven to be the best anti-catabolic treatment thus far.(12,17) Beta-adrenergic receptor blockade using propranolol to attenuate the effect of catecholamines has been proven to reduce the heart rate, lower hypermetabolism, improves immune response, fatty liver infiltration and improve lean muscle mass accretion. (13,17) The use of beta blockers in burn patients has also been associated with decreased mortality, reduce insulin resistance, reduce wound infection rate and wound healing time(12,17)

Good control of glucose has been shown to improve graft take, wound healing, immuno-modulation (improves white cell function, reduces pro-inflammatory cytokines, increases Granulocyte Colony Stimulating Factor [G-CSF] and helps with resistance to infection) and protein balance. Maintenance of blood glucose at or below 110mg/dl helps to reduce mortality.(17) Most of these benefits have been noted but the use of metformin has also been proven to reduce hyperglycemia, reduce muscle catabolism and improve insulin sensitivity without increasing the risk of hypoglycemia.(13) these two agents have a synergistic effect when administered together.(17)

It is estimated that up to 45% of the energy hypermetabolic response is for thermogenesis due to the cooling effect of increased evaporation from burn wounds (up to 4000ml/m^2^ TBSA/day).(11,13) Increasing the ambient temperature to thermo-neutrality (33^11^C) in addition to use of occlusive dressing therefore minimises cooling, thermogenesis and the need for glucose from catabolic pathways.(13)

The finding of increased mortality with fluid administration was largely unexpected. We think that this finding can be explained in two ways. Firstly, the need for intravenous fluid administration denotes the presence or a significantly large burn and/ or shock and these factors are independently associated with mortality. Secondly, there is a possibility of inappropriate fluid administration (either excess or inadequate administration) due to inaccurate assessment of TBSA and or inattentive administration of fluids. Inadequate fluid resuscitation perpetuates shock which is the most common cause of death in the ebb phase (first 48 hours) following a burn.(18,19) there is evidence from multiple studies that adults often get more fluids than predicted by the Parkland formula (4ml/kg/% TBSA) although the underlying mechanisms for such are not fully elucidated.(19–21) The excess fluid often results in increased compartment pressures, oedema, acute respiratory distress syndrome(ARDS), multi organ failure and eventually death.(10,19,21,22) In the absence of an ideal endpoint of fluid resuscitation, careful titration of intravenous fluids according to the parkland formula remains the mainstay of fluid management in our setting(18,23)

To our surprise, patients that presented late to the hospital had a reduced risk of death. We think there are two likely explanations for this. Firstly, these patients had less severe forms of burns and probably had no sense of urgency to come to the hospital. Secondly, QECH is a tertiary hospital and it is possible that patients that had a higher time lapse to presentation were first stabilised at a secondary health care facility.

Our study had several limitations. The data we used in this study was collected as part of routine patient care and record keeping, as such we had a lot of missing data which was excluded in the analyses. This has an effect on the standard errors of our estimates, thereby affecting precision. We categorised our variables in such a way that would minimise data sparsity, hence we did not opt for other advanced ways of dealing with missing data such as multiple imputation Instead of dropping the missing records. Since there were a lot of variables to be tested as risk factors, and there were only 132 primary outcomes of interest, it was decided to limit the number of variables to adjust by using a causal framework formulated in a Directed Acylic Graph (DAG). This is both a strength and a weakness of the study. It is a strength in the sense that our modelling strategy was done using a plausible causal mechanism of death thereby dealing with confounding better than the traditional modelling methods. However since there was no published causal mechanism, we developed our own DAG. Other scientist may disagree with some of the causal paths present or not present.

## CONCLUSIONS AND RECOMMENDATIONS

This is the first ever study in this population and setting to explore the risk factors of mortality in adult burns patients. Important risk factors of mortality include: age, gender, aetiology, burn surface area, burn depth, presence of inhalation burns, circumstances of the burn, quality of co-morbidity management, type of management for the burn wounds, patients residence, time to hospital presentation, length of hospital stay and the type of management the patient gets. In light of our findings, recommend a prospective study in order to avoid the weaknesses of retrospective designs and to explore the impact of the suggested therapeutic approaches (which are currently not practised) on clinical outcomes. We also recommend further studies that explore how factors like age and TBSA modify the effect of the risk factors on outcome of death.

## Footnotes

1 Pro-inflammatory cytokines enhance catabolism and hypermetabolism by inhibition of the growth hormone–IGF-I–insulin axis

